# Convergent evolution of type I antifreeze proteins from four different progenitors in response to global cooling

**DOI:** 10.1101/2024.06.04.597461

**Authors:** Laurie A. Graham, Peter L. Davies

## Abstract

The alanine-rich, alpha-helical type I antifreeze proteins (AFPs) in fishes are thought to have arisen independently in the last 30 ma on at least four occasions. This hypothesis has recently been proven for the flounder and sculpin AFPs that both originated by gene duplication and divergence followed by substantial gene copy number expansion. Here we have examined the origins of the cunner (wrasse) and snailfish (liparid) AFPs. The cunner AFP has arisen by a similar route from the duplication and divergence of a GIMAP gene. The coding region for this AFP stems from an alanine-rich region flanking the GTPase domain of GIMAPa. The AFP gene has remained in the GIMAP gene locus and undergone amplification there along with some GIMAPa genes. The AFP gene originated after the cunner diverged from the common ancestor to the closely related spotty and ballan wrasses that have a similar gene synteny but are completely lacking the AFP genes. Snailfish AFPs have also recently evolved because they are confined to a single genus of this family. In these AFP-producing species the AFP locus does not share any similarity to functional genes. Instead, it is replete with repetitive DNAs and transposons several stretches of which could code for tracts of alanine with a dominant codon (GCC) that matches the bias seen in the AFP genes. All four known instances of type I AFPs occurring in fishes are independent evolutionary events that happened soon after the onset of northern hemisphere Cenozoic glaciation events. Collectively they provide a remarkable example of convergent evolution to one AFP type.

## Introduction

Ice-binding proteins (IBPs) share a common ligand, namely ice, but have a variety of functions including ice anchoring, controlling the growth of ice channels, preventing recrystallization of ice in the frozen state, or preventing internal ice growth in freeze intolerant organisms. They are found in a small percentage of known species, but these are scattered throughout the tree of life and include bacteria, diatoms, insects, plants and fish (1, 2). When they are used to prevent freezing, they are generally called antifreeze proteins (AFPs) and they act by irreversibly binding to ice crystals, lowering their non-equilibrium freezing point (3, 4).

Fish living in ice-laden regions of the ocean often employ AFPs since they freeze at a higher sub-zero temperature than seawater (5). To date, four types of AFP (types I, II, III, IV) and one antifreeze glycoprotein (AFGP) have been described in fish (6). Type III AFP is a globular protein derived from the C-terminal domain of sialic acid synthase (7–9), and it is found exclusively in the infraorder Zoarcales (10). Type II AFP is also globular, but it is derived from a C-type lectin (11, 12). It is found in three separate fish orders that diverged from each other more than 200 Ma and was gained in two of these order through lateral gene transfer (13, 14). Type IV was identified in longhorn sculpin serum and shows similarity to apolipoproteins that form helical bundles (15). However, its serum concentration is insufficient for freeze protection (16) and type I AFPs were subsequently found in the skin of this fish (17), so it may have a function unrelated to freeze protection.

In contrast to the AFPs described above, type I AFP and AFGP are non-globular. AFGPs are extremely repetitive, containing between 4 to ∼50 tripeptide repeats, mostly Ala-Ala-Thr, with an O-linked disaccharide on each Thr residue (18). It adopts a poly-proline type II structure in solution (19) and is found in fishes at both poles. The AFGP gene of Antarctic fishes in the suborder Notothenioidei was derived from a duplicated trypsinogen gene (20). Here the bulk of the gene was lost, but the signal peptide and 3ʹ UTR were retained, along with a 9 bp segment encoding Ala-Ala-Thr that spanned the start of the second exon that was amplified many times. Northern cods from suborder Gadoidei were found to produce similar AFGPs (21), but in this group, they arose from non-coding DNA (22). Type I AFPs are Ala-rich α-helical proteins that are somewhat less repetitive than AFGPs. Many versions have repeats that are 11 a.a. in length, delineated by a single non-glycosylated Thr residue (23, 24). The Ala residues that dominate one side of the helix are well conserved, but residues on the other side are more variable. Like the AFGPs, they vary in length, from 33 to 195 a.a. (24, 25). They are found in three different orders and four different families of fish; the flounders (order Pleuronectiformes) (26–28), cunner (*Tautogolabrus adspersus*, order Labriformes, family Labridae) (29), snailfish (*Liparis atlanticus* and *L. gibbus*, order Perciformes, family Liparidae) (30) as well as a number of sculpins, including the shorthorn sculpins (*Myoxocephalus scorpius*, family Cottidae) (17, 24, 31, 32) (Fig. 1). The shorter type I variants are less active and form isolated helices, often with N- and C-terminal modifications (33, 34), whereas the longest, hyperactive variant (Maxi) folds in half and dimerizes to form a four-helix bundle stabilized by internal waters (35).

**Figure 1:**
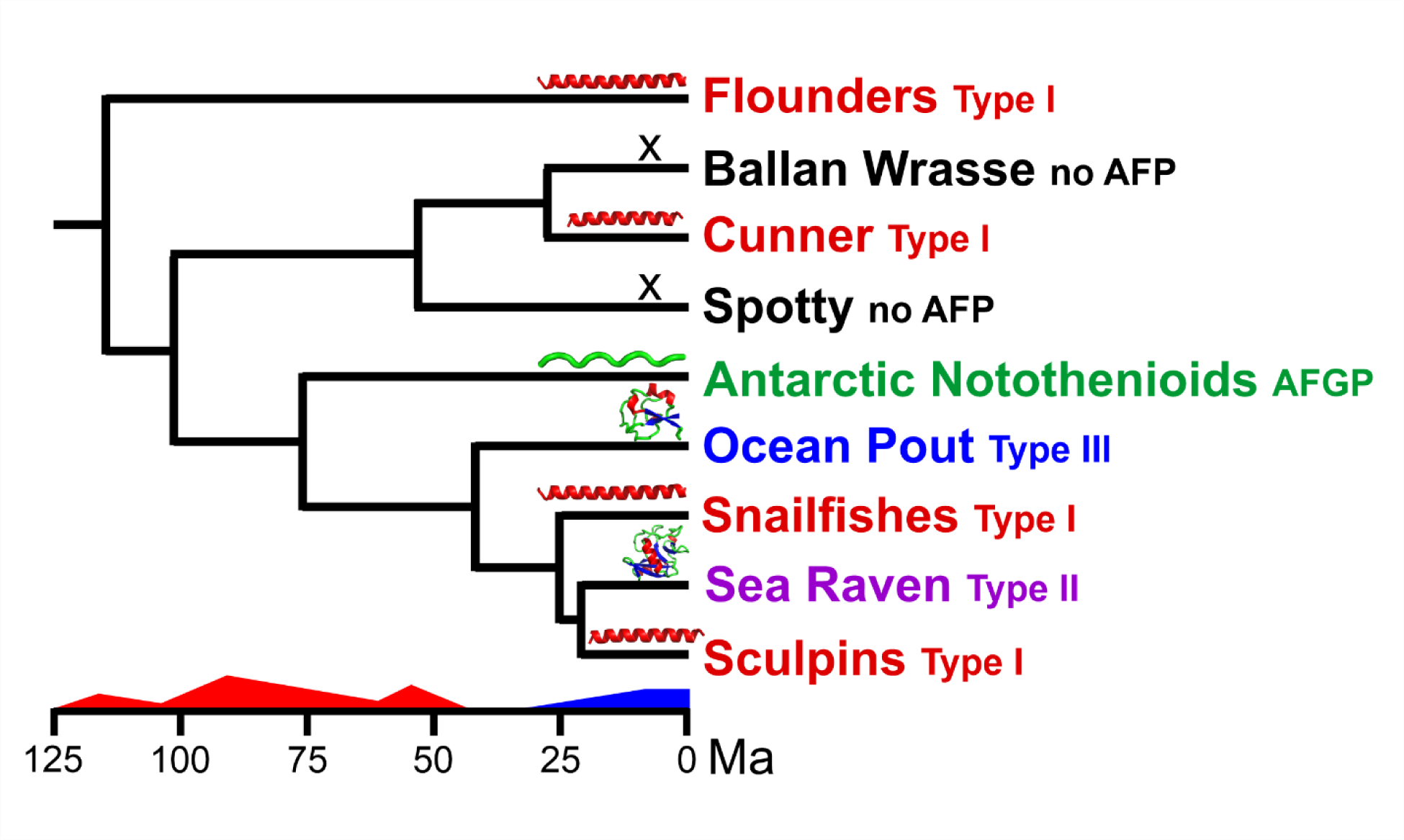
Divergence of AFP-producing fishes during changing climatic conditions. The relationships between the fishes or fish groups shown, all within in clade Percomorphaceae, were obtained from a time-calibrated phylogeny of almost 2000 fishes (79) or over 200 Labrids (80). While the majority of the species within this clade do not produce AFPs, only those examined in this study are included (black font). Cartoon graphics of the AFP types generated using PyMOL (81) are shown along each branch. The timing and intensity of the warmest (red) and coldest (blue) climactic periods of the last 125 Ma are indicated above the time scale (78).

Type I AFPs may have pro-peptides and/or signal peptides, or they may have neither. All three variations are found in flounder AFPs (25, 26, 36), whereas cunner, sculpin and snailfish AFPs lack signal peptides and are still secreted (17, 29, 30, 32). While most of these AFPs have Thr-residues at 11 a.a. intervals, those from snailfish (30), and one from shorthorn sculpin (37), do not. Interestingly, the skin isoforms of flounder are more similar to sculpin sequences than they are to either the liver or maxi isoforms. However, the patchy taxonomic distribution of type I-producing fishes (Fig. 1), lack of similarity between the untranslated regions (UTRs) and differential codon usage led us to suggest that these proteins were not homologous and that their similarities arose through convergence (38).

Phenotypic similarities frequently arise independently, often but not exclusively, when similar environmental challenges are encountered, and it has been argued by Michael P. Speed and Kevin Arbuckle that “analyses of convergence should typically be paired with broader investigations of the evolutionary history of the trait” (39). This has now been rectified for type I AFPs as the last piece of the puzzle necessary to unequivocally demonstrate their convergence in these four fish groups, namely a demonstration of how they arose, has been achieved for all of the lineages. We recently determined that the flounder *AFP* arose from *Gig2* (40), which encodes a protein involved in viral resistance (41). Subsequently, we found that the sculpin AFP arose from *lunapark* (42), a gene whose protein resides within the endoplasmic reticulum and stabilizes membrane junctions in this organelle (43). Here, we demonstrate that the cunner AFP also arose from a different pre-existing gene, confirming its origin via convergent evolution. The snailfish *AFP* does not resemble any gene loci. Instead, the coding sequence is tightly flanked by transposons from which it likely originated.

## Materials and Methods

### BLAST searches and databases

Known type I AFP sequences (nucleotide and protein) were used as queries at the BLAST interface of NCBI (44), limiting the taxa searched to Teleost fishes. Low complexity filters, masks and/or compositional adjustments were turned off and moderate stringency was selected (discontiguous megablast, blastn, or blastp with BLOSUM45 matrix) to detect more divergent sequences. High-throughput genomic and transcriptomic sequence datasets were accessed through the NCBI SRA portal (https://www.ncbi.nlm.nih.gov/sra) and genome assemblies through the genome portal (https://www.ncbi.nlm.nih.gov/genome/).

### Analysis of cunner AFP sequences

A 250-kb segment of chromosome 4 that contained AFP genes was downloaded from NCBI (GenBank JAJGRF010000003.1, 6,121,000 to 6,371,000 bp) from the representative genome of cunner (GCA_020745675.1). Gene annotation was done in SnapGene Viewer (from Insightful Science; available at snapgene.com) using BLASTn to identify AFP coding sequences and GeneWise (45) to identify AFP-adjacent genes based on homologues in the annotated genomes of the closely-related ballan wrasse (*Labrus bergylta* GCA_900080235.1, scaffold NW_018114954.1, 83944 to 184,596 bp) and New Zealand spotty (*Notolabrus celidotus* GCA_009762535.1, chromosome 7, NC_048278.1, 5,367,202 to 5,491,088 bp). Data use policy: https://genome10k.ucsc.edu/data-use-policies/

### Phylogenetic comparisons of GIMAP sequences

The structure of the *GIMAP* genes of the ballan wrasse were verified or reannotated based upon matching reads from transcriptomic sequences (Accession SRR5454465) from the NCBI Sequence Reads Archive (SRA). The spotty *GIMAP* genes were reannotated, when necessary, by comparison with teleost GIMAPs from the non-redundant (nr) protein database. The sequences of the GTPase domain of the GIMAPs of the cunner, spotty and ballan wrasses were aligned in SeaView version 5.0.1 (46) and are shown in Supplementary Fig. 1. A maximum likelihood phylogenetic tree was generated in MEGA 10.1.8 (47) using model JTT G+I with 100 bootstrap replicates.

### Acquisition of an elongated genomic sequences containing a snailfish AFP gene

A DNA sequence closely matching the known dusky snailfish AFP (30) was selected as an assembly seed from the genomic DNA SRA dataset (GenBank SRR22396815). As the dataset was large, and provided good coverage, overlapping reads that matched with 100% identity over 80% of their length were iteratively added to the growing assembly using GeneStudio V2.2.0.0. (GeneStudio Inc.).

### Derivation of codon usage statistics

The frequency at which Ala codons were used in various sequences was determined using the online Codon Usage Calculator from Biologics International Corp (Indianapolis, IN, USA https://www.biologicscorp.com/tools/CodonUsageCalculator/). The sequences used for cunner were the combined coding sequences of the 11 AFP isoforms as well as the segment encoding the Ala-rich C-terminal region of GIMAP-a5 (56 a.a.). For the ballan wrasse, the sequence encoding the last 114 a.a. of GIMAP-a1 was used. For snailfish, the nine sequences encoding the AFPs shown in Fig. 5 were used. Ala codon usage in Teleost fishes was taken from the CoCoPUTs database (48), in which 5,376,783 Teleost coding sequences had been analyzed as of March 3, 2022 (https://dnahive.fda.gov/dna.cgi?cmd=codon_usage&id=537&mode=cocoputs).

**Figure 2:**
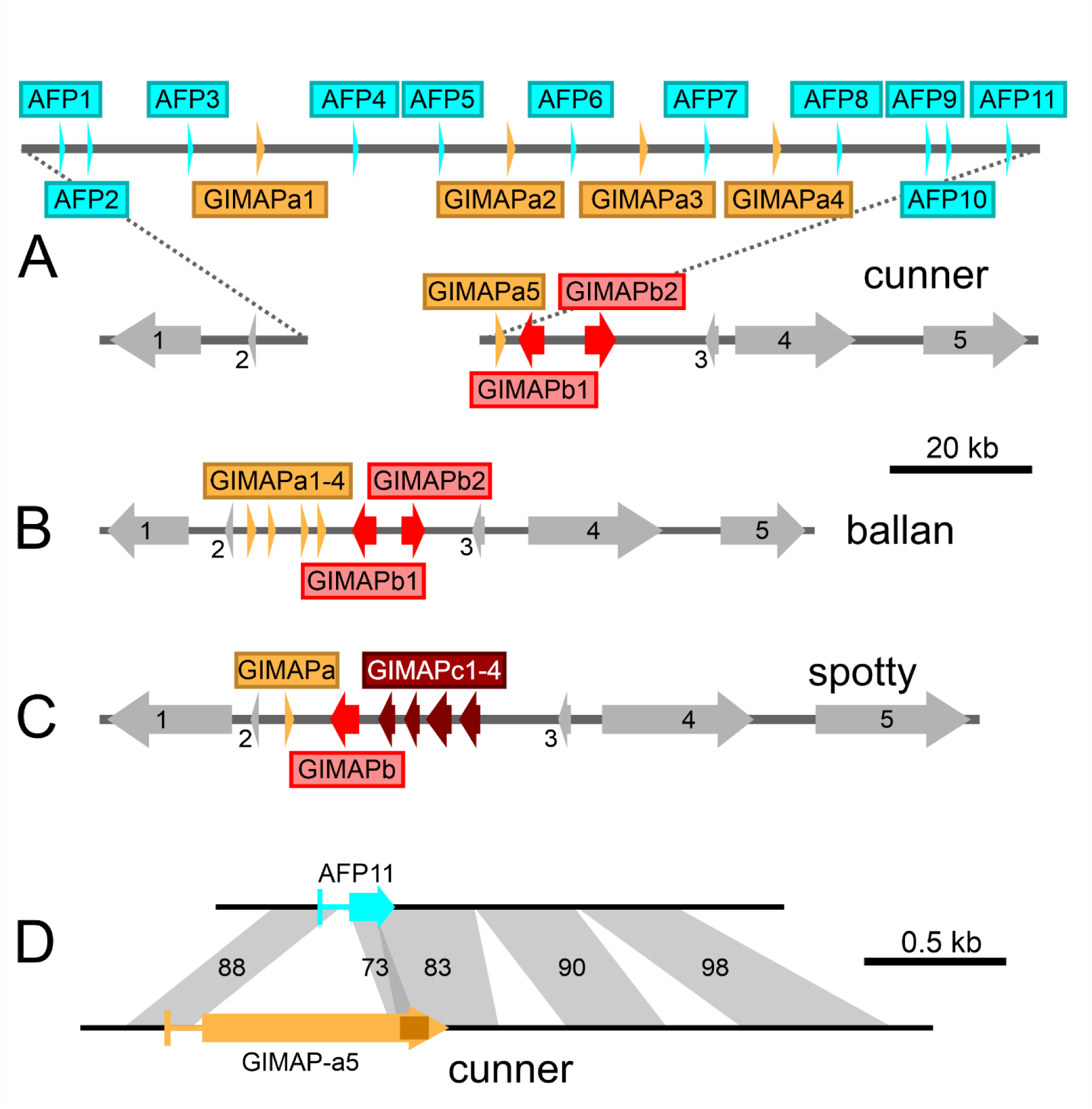
*AFP* locus in cunner compared to the syntenic location in two other wrasses. A) Cunner AFP locus on chromosome 4, showing locations of the 11 *AFP* genes (cyan) relative to the *GIMAPa* genes (dark yellow) and *GIMAPb* genes (red). The arrows indicate gene orientation (all *AFP* genes and *GIMAPa* genes are transcribed left to right) and span the coding region and intron(s). Flanking genes are numbered consecutively in grey. The scale bar shows a 20-kb stretch of DNA. B) Ballan wrasse locus coloured as above. C) Spotty locus, coloured as above with *GIMAPc* genes in dark red. The unrelated flanking genes, numbered sequentially, are solute carrier family 25 member 10-like (SLC25A10) and claudin-9-like (CLDN9) at the 5ʹ side, ras-related dexamethasone-induced 1-like (RASD1), MYCBP associated protein-like (MYCBPAP) and epsin 3-like (EPN3) at the 3ʹ side. D) Comparison of the cunner *AFP11* and *GIMAP-a5* genes. The percent identity between the homologous regions (grey shading) is indicated, with the two exons (wide) and intron (narrow) of each gene in cyan or dark yellow. The portion of the second exon of GIMAP-a5 that contains an Ala-rich region is in a darker shade. The scale bar shows a 0.5-kb stretch of DNA.

**Figure 3:**
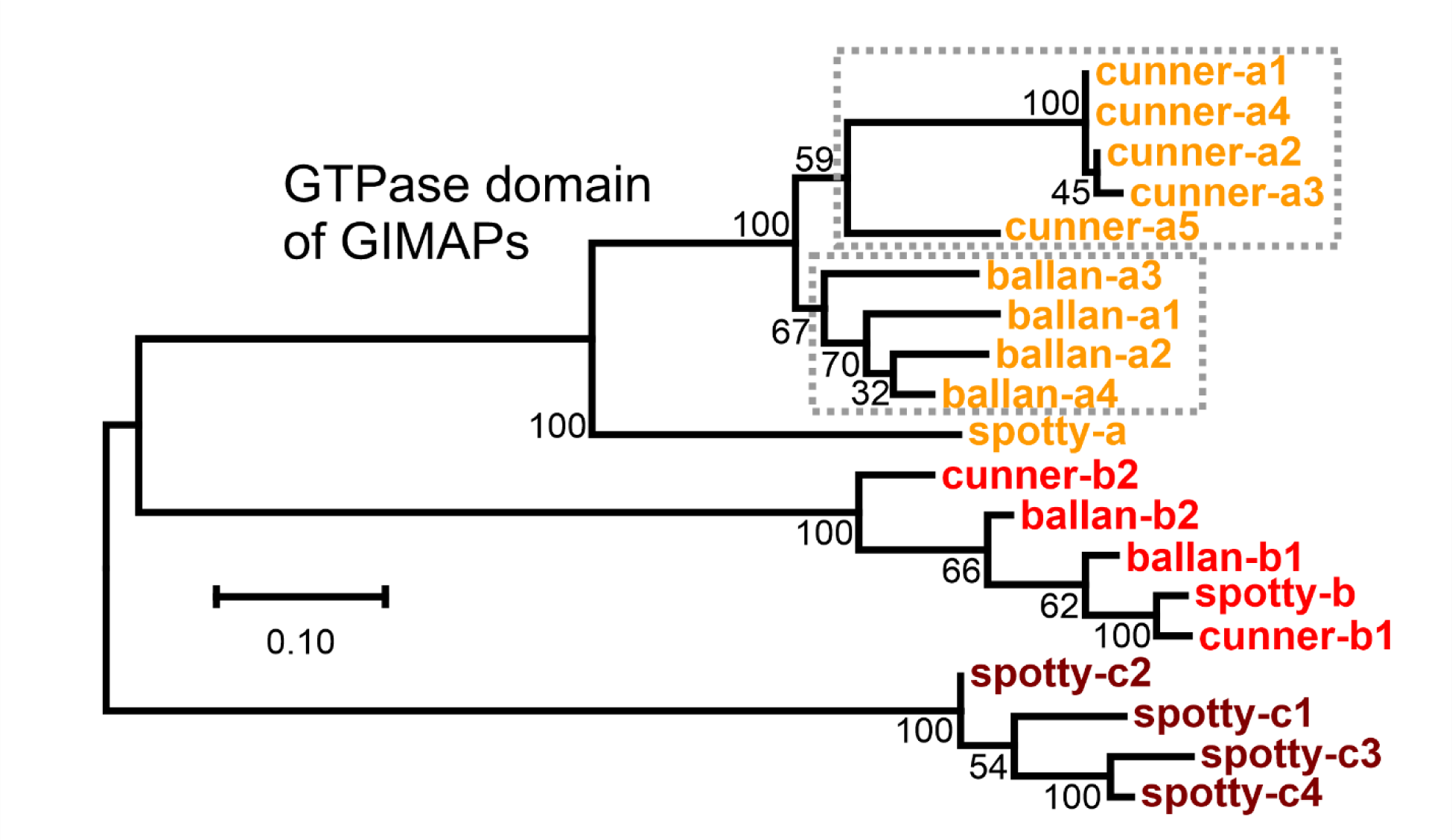
Phylogenetic relationships of GIMAP proteins found in the three wrasses from maximum-likelihood analysis of an alignment of the GTPase domains (Supplementary Fig. 1). The coloring of the labels matches the coloring of the genes in Fig. 2. The bootstrap values (%) are shown at each node. Note that cunner-a1 and -a4 are identical.

**Figure 4:**
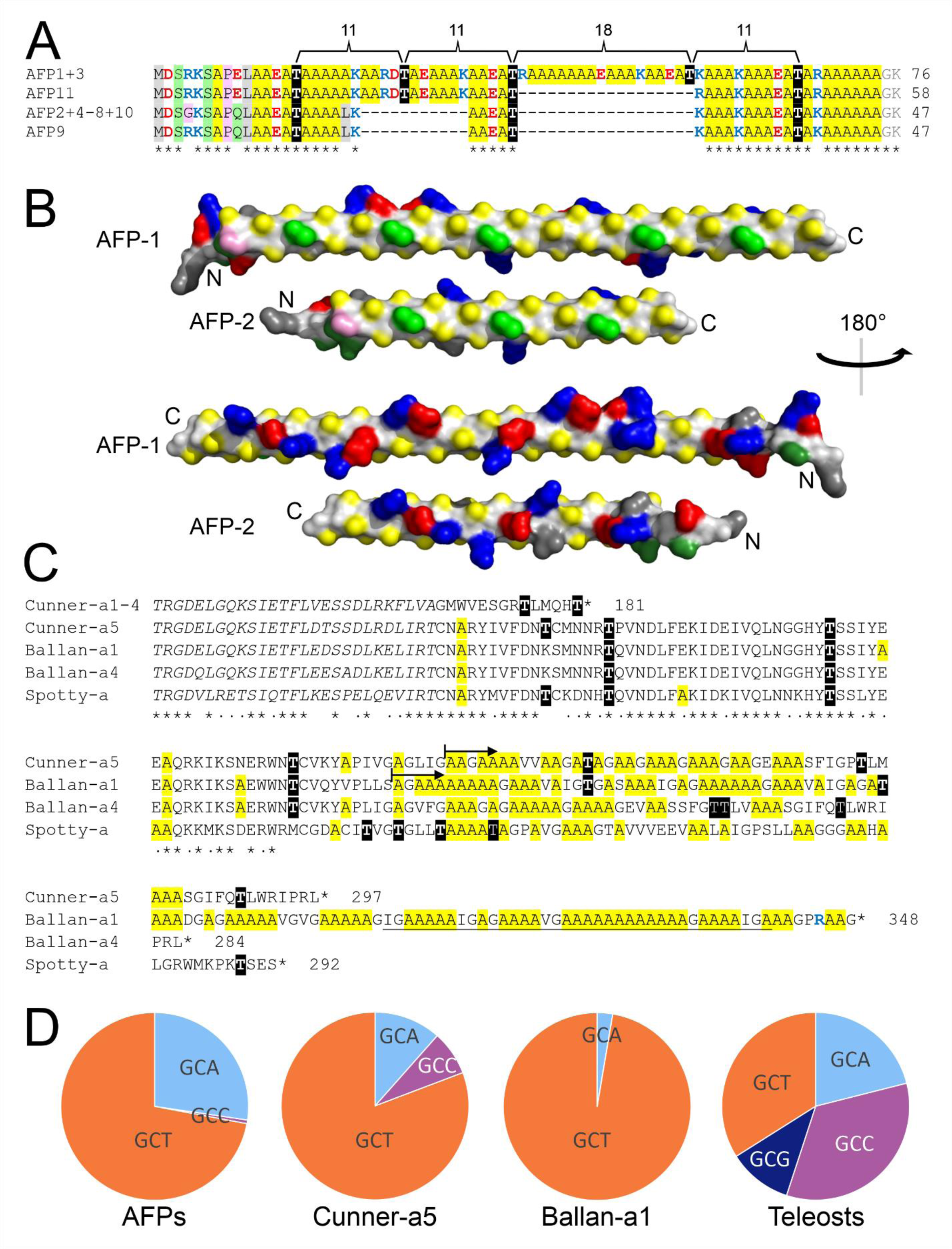
Sequences, models, and codon usage of cunner AFPs and Ala-rich C-terminal regions of GIMAP genes. A) AFP isoforms aligned with Ala highlighted in yellow, acidic and basic residues in red and blue font, respectively, Gly and Pro highlighted pink, polar residues other than Thr highlighted green, aliphatic residues highlighted grey, with the spacing of Thr residues (black highlighting) indicated above and asterisks indicating 100% conservation below. The last two residues (faded grey) are naturally removed when the C terminus is amidated (29). B) Models of the long (AFP-1) and short (AFP-2) cunner AFPs generated using AlphaFold2-Colab (82), rendered using PyMOL (81). Residues are coloured as above but with backbone atoms in light grey, Thr in green and other polar non-charged residues in dark green. Two 180°-degree rotations are displayed with their termini marked N and C. C) Ala-rich C-terminal regions of Cunner-A5 and Ballan-A1 GIMAPs relative to the shorter Cunner A isoforms. Ala and Thr residues within the extension are coloured as in A above with the end of the GTPase domain in italics. Conservation between the unambiguously aligned residues of the four isoforms with extensions is indicated below the alignment by asterisks (all four sequences identical) or dots (three of four sequences identical). The beginning of the two segments used to derive Ala codon usage are indicated with arrows. D) Ala-codon usage of cunner AFPs (all isoforms, 322 codons) compared to the Ala-rich extensions of Cunner-A5 (26 codons), Ballan-A1 (75 codons), and Ala codons sampled over 5 million coding sequences from teleost fishes.

**Figure 5:**
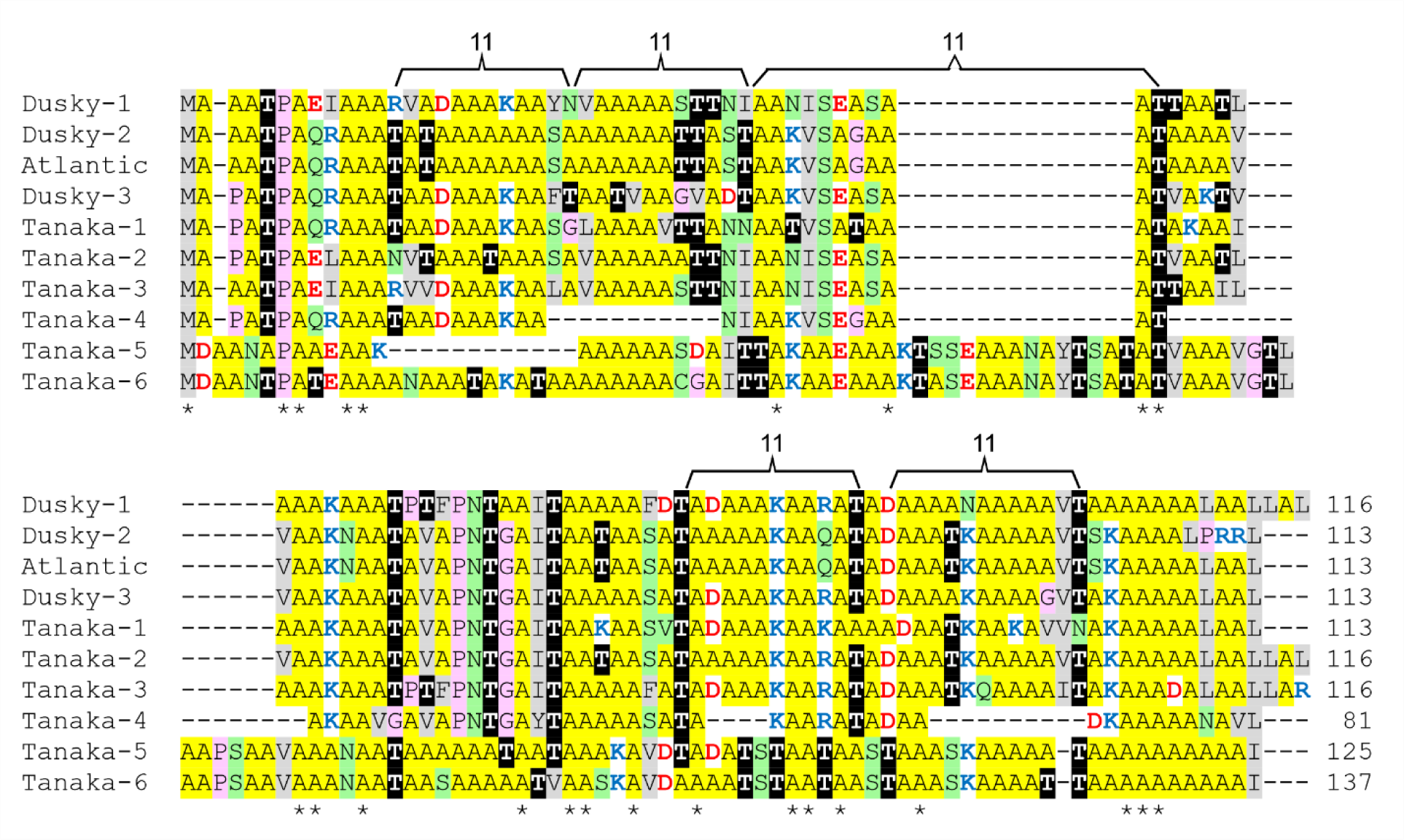
Alignment of snailfish AFP sequences coloured and annotated as in Fig 4A. The GenBank accession numbers of the DNA sequences encoding these isoforms are Dusky-1 from a transcriptome, MT678484; Dusky-2 from a cDNA, AY455863; Atlantic from a cDNA, AY455862; Dusky-3 assembled from genomic SRA sequences (Supplementary Fig. 3); Tanaka-1 to −5 from genomic DNAs, SRLO01012854, SRLO01001937, SRLO01021987, SRLO01024077, SRLO01015817 respectively; Tanaka-6 from bases 3,853,802 to 3,857,801 of CM070098.

## Results

### Part 1: Cunner

#### Cunner AFPs reside at a single locus

The cunner reference genome, included in the Vertebrate Genomes Project (49), was screened for AFP sequences using the cunner cDNA sequence (29). Matches were found at a single location, spanning 133 kbp, on chromosome 4 (Fig. 2A). This genome was not annotated, so the eleven *AFP* genes found here were annotated based upon the known cDNA sequence (Fig. 2A, cyan arrowheads). Additionally, seven interspersed (yellow and red arrowheads) and five flanking genes (grey arrowheads) in the immediate neighbourhood were also identified and marked based upon the annotated genomes of the spotty and ballan wrasses (49), two closely-related fishes in the same family (Labridae, commonly called wrasses). The microsynteny of the flanking genes is conserved between the three species (Fig. 2, grey arrows), with the five encoded proteins sharing 89 – 99% identity between cunner and ballan wrasse. As expected, the identities between cunner/ballan wrasse and the more distantly-related spotty are lower, ranging from 73 – 95%. The two other cunner assemblies (pseudohaplotype GCA_020745675.1 and GCA_024362835.1) were incomplete in this region, underscoring the difficulty of assembling multigene families.

#### AFP genes in the cunner are interspersed with GIMAP genes and share sequence similarity

A total of seven proteins belonging to the GTPase IMAP family (GIMAPs) were encoded by genes found interspersed amongst the *AFP* genes of the cunner (Fig. 2A). The ballan wrasse and spotty each had six *GIMAP*s (Fig 2B-C), but *AFP* genes were not present at these loci, nor elsewhere in these genomes. The *AFP* genes share both proximity and sequence similarity with the *GIMAP* genes. The pairing with the highest similarity is *AFP11* with *GIMAP-a5*, where five segments have identities ranging from 73 to 98% (Fig. 2D). These segments lie both upstream and downstream of the coding sequence and overlap the first exon and the majority of the intron. The most notable difference between the loci is that the majority of the coding sequence within exon 2 is absent from the AFP. These similarities are sufficient to indicate that the AFP gene arose from a duplicated *GIMAP-a* gene.

Before the GIMAP genes were compared between the cunner, spotty and ballan wrasse, errors in the automated annotation of this repetitive gene family were corrected as described in Materials and Methods. The accession numbers and sequence, if modified, are shown in Supplementary Table 1. Phylogenetic analysis indicated that the GIMAPs of these three species cluster into three groups, herein labelled as type a, b or c (Fig. 3). Type c is restricted to spotty (four isoforms) where there is just one each of the type a and type b isoforms. Ballan wrasse and cunner have two divergently transcribed type b genes and four or five type a genes. These type a proteins cluster along species lines, with shorter branch lengths between the cunner isoforms, indicating that these genes were duplicated after the divergence of these two lineages and that this occurred more recently in the cunner. Taken together, this indicates that the *GIMAP* gene family is dynamic and that the AFP genes arose from a *GIMAP-a* gene, with both genes being subsequently amplified in tandem within the cunner lineage.

#### The eleven cunner AFPs are highly similar

There are four AFP isoforms encoded by the eleven *AFP* genes (Fig. 4A), seven of which (2, 4–8, 10) match the previously characterized sequence (29). Over 50% of the residues are Ala, and with one exception, Thr is spaced at 11-residue intervals. AFP9 differs from the main sequence at a single position (residue 4, Gly to Arg) while AFP11 contains one additional 11-a.a. repeat. Two genes (*AFP1* and *AFP3*) encode identical isoforms that match AFP11, except that they contain an 18-a.a. insertion in which the Thr residues are spaced 18 residues apart.

AlphaFold2 models of both the longest (AFP-1) and shortest (AFP-2) isoforms, in which the longer isoform has 29 additional residues, are very similar (Fig. 4B). Both form extended amphipathic α-helices that have a hydrophobic, Ala-rich surface punctuated by Thr residues. The other side of the helix is also enriched for Ala, but all of the charged residues are found here, many of which appear to form helix-stabilizing salt bridges. The disruption in the 11 a.a. spacing of the Thr residues by one 18 a.a. segment in the long isoform is an interesting deviation. An exact periodicity of 11 a.a. would correspond to 11 residues/3 turns, or 3.67 residues/turn, whereas the typical α-helix has 18 residues/5 turns, or 3.60 residues/turn. An examination of winter flounder AFP isoforms, including the crystal structure of a short isoform (34) and NMR structure of an engineered variant (50), as well as the crystal structure of the hyperactive isoform (35), reveal that residues at 11 a.a. intervals have a slight precession as the periodicity is closer to ∼3.65 residues/turn. Therefore, the 18 a.a. insertion serves to counteract this, bringing the Thr back into register (Fig. 4B, top).

#### Cunner AFP arose from the C-terminus of the GIMAP-a protein

The Ala content of the GIMAP proteins is generally low. For example, cunner GIMAPa-1 has only seven Ala residues, making up 4% of the total. The four exceptions to this are cunner GIMAPa-5 (12%), ballan wrasse GIMAPa-1 (24%) and GIMAPa-4 (11%), plus spotty GIMAPa (10%). Their Ala-richness is restricted to the C-terminal region, outside of the GTPase domain, as shown in Fig. 4C. These extensions are present in all three wrasses being compared, whereas AFPs are found only in the cunner, so the Ala-rich extension arose prior to the AFP.

While these extensions are rich in Ala, they lack the periodicity of the Thr residues and contain more Gly, and fewer charged residues, than the AFPs. An AlphaFold2 model of the isoform with the longest C-terminal extension (Ballan-a1, Fig. 4C) predicts three α-helical segments in this region (Supplementary Fig. 2A), with the last spanning 37 aa (underlined in Fig. 4C) with 27 Ala residues (73%). The Ala-rich region of the shorter extension of Cunner-a5 is predicted to be unstructured (Supplementary Fig. 2B). Nevertheless, there is sufficient sequence similarity (Fig. 2D, darker yellow) to indicate that the Ala-rich extension gave rise to the AFP. The partial overlap of two of the matches is consistent with an internal duplication within the longer AFP11 allele. Interestingly, the similarity between the coding sequences is lower than in non-coding regions, consistent with positive selection of the AFP for its new function. Taken together, these data indicate that the AFP arose from a duplicated *GIMAP-a* gene containing an Ala-rich extension from which the GTPase domain was lost.

#### Ala codon usage of GIMAP-a and AFPs is similarly atypical

A further line of evidence that supports the Ala-rich extension of GIMAP-a as being the progenitor of the AFP is that they share a similar codon usage bias. Cunner AFPs are unique amongst the type I AFPs in that Ala is preferentially encoded by GCT (72%), rarely by GCC (<1%), and not at all by GCG (Fig. 4D). In contrast, in teleost fishes, GCT and GCC each encode about a third of all Ala residues, while GCG encodes 11%. A similar bias is observed in the Ala-rich extensions, with Ballan-a1 employing GCT almost exclusively. This GCT bias is not observed in the flanking genes of any of these fish (not shown), indicating that it is a characteristic of the C-terminal extension of the *GIMAP-a* genes that was retained in the *AFP* genes.

### Part 2: Snailfish

#### AFP sequences are only present in one genera of snailfishes

BLAST searches of genome sequences, the transcriptome shotgun assembly, and a selection of SRA datasets using both cDNA and protein sequences from snailfish AFPs (30, 51) shows that in addition to species previously known to produce AFPs, namely Atlantic (*Liparis atlanticus*), dusky (*L. gibbus*) and Tanaka’s (*L. tanakae*) snailfish, they are also found in *L. liparis* and *L. tunicatus*. Similar searches failed to identify homologues in other members of the same family (Liparidae) in different genera (Supplementary Table 2). However, the low complexity of the snailfish AFPs (55-61% Ala, encoded primarily by GCC), means that divergent AFP sequences would be difficult to identify.

#### Snailfish AFPs are members of a multigene family

Tanaka’s snailfish was the only member of this genus with assembled genomes, one consisting of almost 28,000 scaffolds (52) and the second with 24 chromosomes and 902 unplaced scaffolds (53). Three AFP loci had been previously identified (51), but at least six additional loci are present, five of which were obtained from the first assembly, but only one from the second (Supplementary Table 3). All but one of the scaffolds are under 3 kb in length, and the longest (25 kb, SRLO01001937) has seven gaps. For three of the scaffolds, gaps are found within the coding sequence.

#### A dusky snailfish locus was assembled from SRA reads

There are several cDNA sequences known for snailfish AFPs, but only two have both 5ʹ and 3ʹ untranslated regions, one from the dusky snailfish (*Liparis gibbus*) (30) and one from the Atlantic snailfish (*Liparis atlanticus*) (54). However, the 5ʹ UTR is quite short (27 bp). To facilitate gene comparisons between species, a longer sequence (1.1 kb, Supplementary Fig. 3) was carefully assembled from a dusky snailfish genomic SRA database, using only reads with significant overlap and 100% identity. The contig contains over 400 bp of upstream sequence and it encodes Dusky-3 (Fig. 5).

#### Snailfish AFPs vary in length and sequence and largely lack regular Thr periodicity

An alignment of the ten AFP sequences reveals three size classes, with seven sequences ranging from 113-116 a.a. in length (Fig. 5). Within this group, Atlantic and Dusky-3 are almost identical, with four differences restricted to the C terminus. The others share 65 to 89% sequence identity. The other three, Tanaka-4, −5 and −6, are more divergent (∼50% identity) and vary in length and sequence, with insertions and deletions relative to each other and to the other sequences.

The 11-a.a. Thr periodicity, prevalent in the cunner AFPs (Fig.4A), is largely lacking in the snailfish AFPs (Fig. 5). The only sequence with four Thr showing this spacing is Dusky 3. All other sequences have zero to two pairs of Thr residues with this spacing. Nevertheless, Thr is the second most abundant residue in all of these sequences, ranging from 9 to 14%. Like all Type I AFPs, Ala is the dominant residue, ranging from 53 to 61%. Another notable difference is the snailfish sequences all have two or more helix-breaking residues (Gly or Pro) around their midpoints (Fig. 5) that are lacking in cunner (Fig. 4A).

#### The gene loci of the Tanaka and dusky snailfishes share similar TEs in their flanking regions

The assembled contigs contained enough flanking sequences to permit detailed comparisons between the loci (Fig. 6A). The assembled dusky gene (dusky-3) is very similar to one of the Tanaka genes (Tanaka-1) and may be an ortholog. It also suggests that both assemblies, obtained independently and by different methods, are accurate. The rest are likely to be paralogs. All of the Takana coding sequences were verified as they each exactly matched many overlapping genomic SRA reads with 100% identity. None of the genes were a 100% match to reads within the one conspecific transcriptome available, but some were close (Supplementary Table 3). Comparisons between these genes and the known cDNAs indicate that these genes do not contain introns.

**Figure 6:**
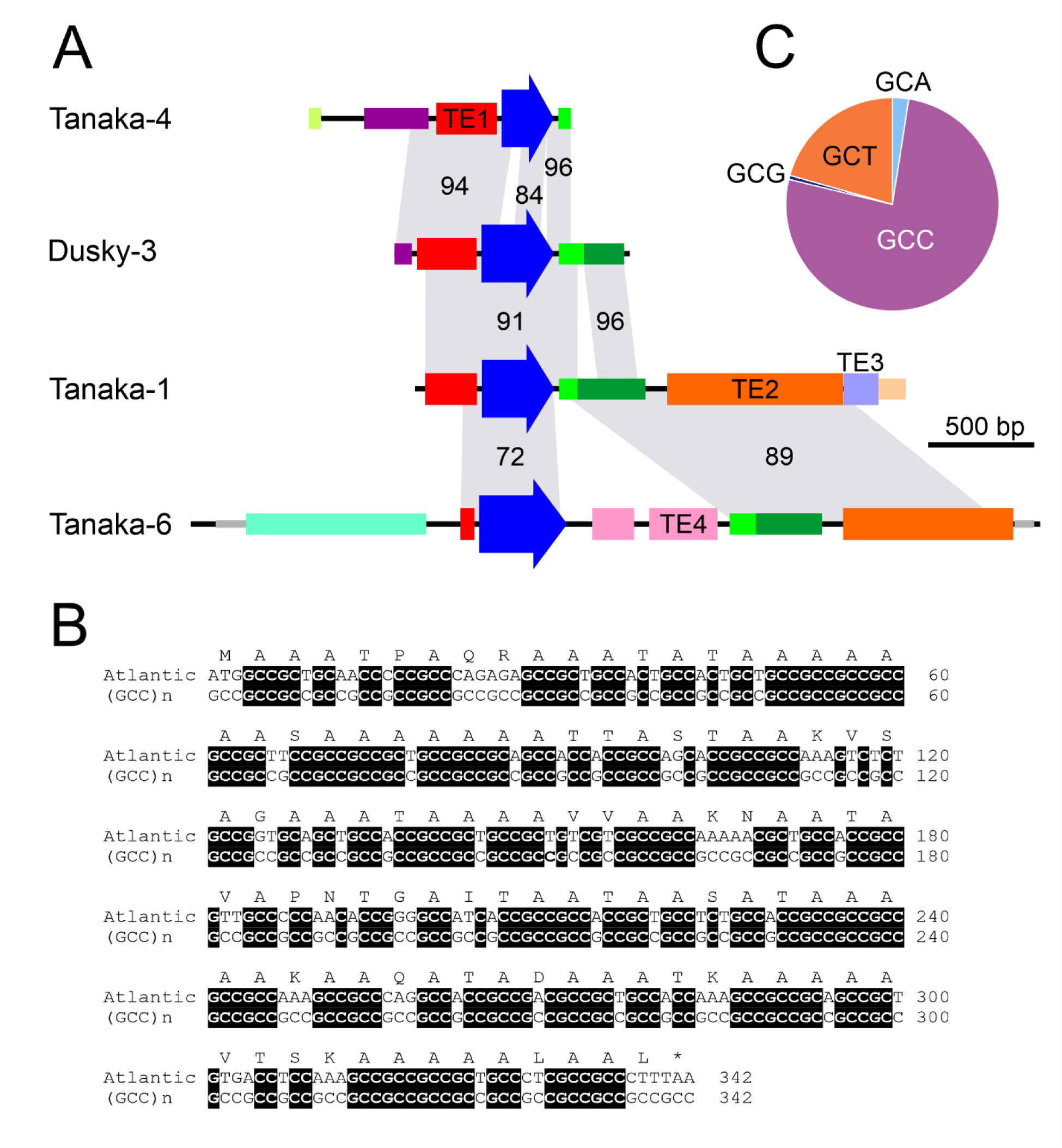
A comparison of snailfish AFP genes with respect to repeats, sequence identities, and codon biases. A) Schematics of four genes encoding AFPs from Fig. 5. The coding sequence is indicated with a dark blue arrow, repeats identified as transposable elements (TEs) by Censor (55) are indicated with wider bars and coloured by type, other repetitive sequences are indicated by narrower bars with colour indicating similarity, and simple repeats are shown with very narrow grey bars. Matching segments are indicated with grey shading with percent identity indicated. B) Alignment of Atlantic snailfish *AFP* coding sequence to a GCC repeat sequence. C) Ala codon usage in the AFPs from Fig. 5.

The flanking regions of all loci lack similarity to known genes. Rather, they are dominated by repetitive DNAs. Some of these segments are similar to transposable elements (TEs) detected by Censor (55), shown as wide bars in Fig. 6A and listed in Supplementary Table 4. Additional segments (thinner bars) showed over 80% identity to sequences present at least 100 times in several different fish genomes, suggesting they are likely TEs that are not present in the Censor database. All of the coding sequences are flanked by the same two TEs, provided they extend far enough. The exceptions are the cDNAs which are too short at their 5ʹ ends (Dusky-1 and −2, Atlantic), and Dusky-2 and Tanaka-5 that are too short at their 3ʹ ends (not shown). The length of the 5ʹ TE varies between loci, but it is always just upstream of the coding sequence. The downstream TE (green) is usually just downstream of the gene, except in Tanaka-6 where two additional TE segments intervene. Interestingly, the identities between the flanking regions that overlap is higher than with the coding sequences (Fig. 6A), a possible indication of positive selection on the AFP sequences.

#### Snailfish AFP likely arose from non-coding DNA and transposons

Given that the flanking sequences of the flounder, cunner, and sculpin genes were clearly associated with progenitor genes, it was presumed that the same would be the case for snailfish. This may well be the case, although rather than being associated with any one gene, they are associated with a variety of TEs and putative TEs, suggesting that these were the progenitors (Fig. 5A). Additionally, there are many instances of simple repeats within Liparid genomes that have the potential to encode runs of Ala residues (not shown). Other fishes have transposons with repetitive segments, such as the Siamese fighting fish (*Betta splendens*) whose Dada transposon (56) contains several stretches with Ala coding potential, the longest of which is 177 bp. An extreme example is (GCC)_151_, found within the last intron of the gene encoding potassium voltage-gated channel subfamily H member 5 (GenBank: XP_048106505) of the Allis shad. Interestingly, the Atlantic snailfish coding sequence can be aligned to these repeats with 72% identity (Fig. 5B). Such an origin would explain the biased codon usage (Fig. 6C) in which 76% of Ala codons are GCC, in contrast with cunner AFP, where GCT is dominant (Fig. 4C). Additionally, these AFP genes lack similarity to any known genes in two other members of the infraorder Cottales with robust assemblies, *Taurulus bubalis* (GenBank GCA_910589615.1) or *Cyclopterus lumpus* (GCF_009769545.1), despite TE1-like sequences sharing over 90% identity between these three species.

#### The AFPs of snailfish form a folded helix and may be dimeric

Snailfish AFPs are known to be largely helical (57), but there are two short regions where this is likely not the case. At the N-terminus, there are one or two Pro residues that may prevent this segment from forming a helix (Fig. 5). The rest of the protein is roughly bisected by two helix-breaking residues spaced three to five residues apart. This is reminiscent of the large isoform of winter flounder where a 195-a.a. alpha helix folds in half and then associates with another molecule as antiparallel dimers that forms a four-helix bundle (35). Therefore, the snailfish AFP was modelled here both as a monomer and a dimer.

The models that were generated (Fig. 7), whether for the monomer or the dimer, folded the polypeptide chain in the same manner. The first five residues were not predicted to form part of the helix. The rest of the chain was helical, with the exception of the bend, punctuated by Pro and or Gly residues (pink). The helical segments on either side of the bend are predicted to lie alongside each other. In the dimeric model, the two monomers are antiparallel.

**Figure 7:**
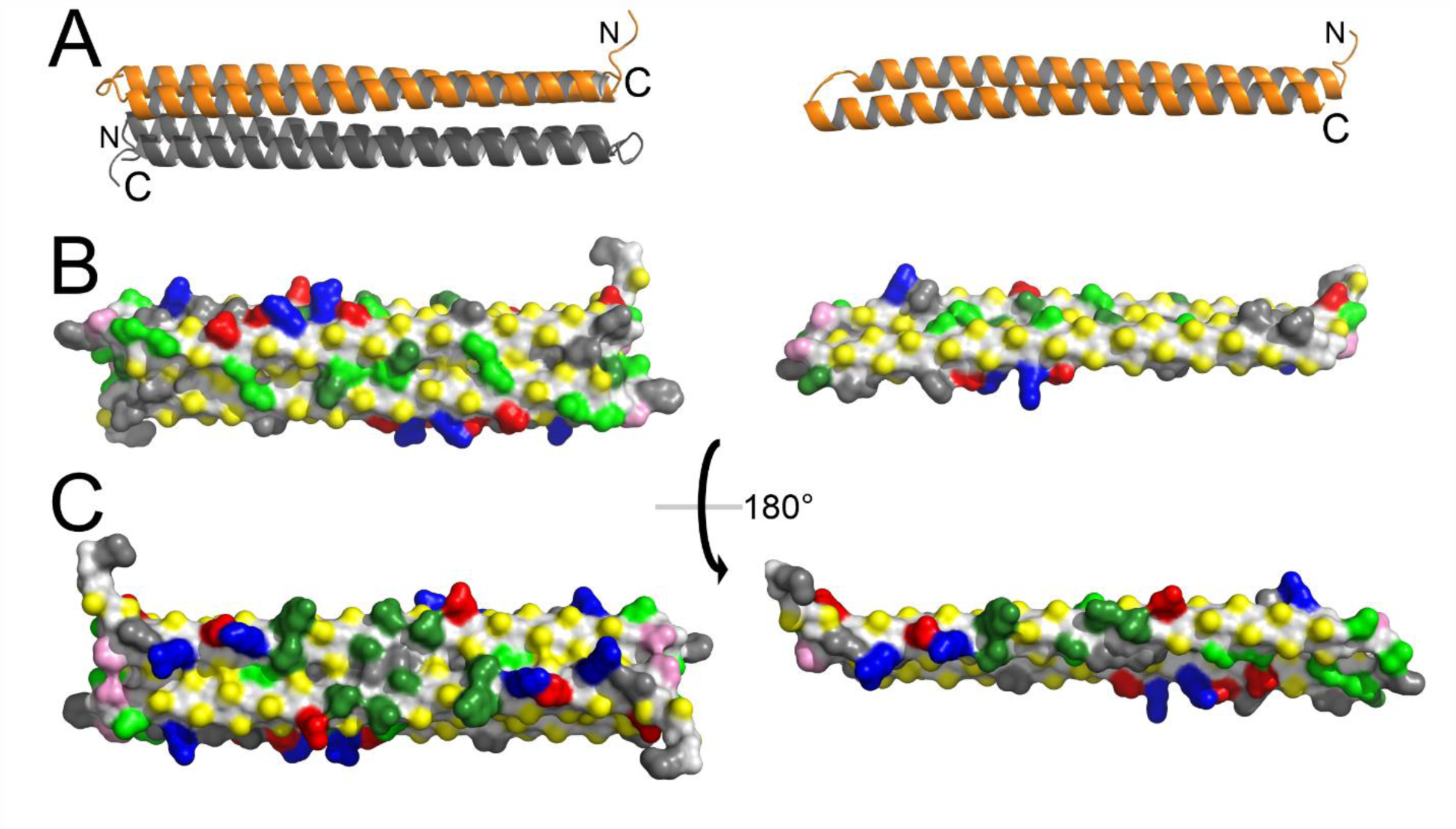
Models of dusky-1 AFP as a dimer (left) and monomer (right) generated using AlphaFold2-Colab (82), rendered using PyMOL (81). A) Cartoon representation of helices with N (back) and C (front) termini indicated. B) Space filling model with the putative ice-binding surface facing forward. Residues are coloured as in Fig. 4 but with backbone atoms in light grey, Thr in green, and other polar residues in dark green. C) View of the reverse side relative to B.

Interestingly, the surface of once side of these models is very flat (Fig. 7B), and like other type I AFPs, it is dominated by Ala and Thr and devoid of charged residues. The dimer also appears plausible as there are several potential intermolecular salt bridges predicted from the antiparallel pairing (red and blue).

### Part 3: Summary of the convergent origins of the four type I AFPs

The AFPs of flounders, sculpins, cunner and snailfishes arose recently enough, sometime within the last 30 Ma (Fig. 1), that their origins could be traced due to the similarities that their non-coding regions have to other sequences. The origins of three of the four type I AFPs were traced to functional genes that were duplicated and underwent divergence. The cunner *AFP* arose from the GIMAP-a gene (Fig. 2), whereas the flounder *AFP* arose from a Gig-2 gene (40), and the sculpin *AFP* arose from a lunapark gene (Fig. 8) (42). In the flounder, the antiviral Gig-2 genes were duplicated at a new location (not shown), and the AFP gene arose from a single copy of the pre-existing Gig-2 gene (Fig. 8B). It was later duplicated at the site of origin multiple times, giving rise to a single locus containing multiple AFP genes in tandem (Fig. 8A), but Gig-2 genes were not retained at this location. In cunner (Fig. 2D), as in flounder (Fig. 8B), the gene structure and much of the flanking sequence of the progenitor was retained. However, in cunner, the GIMAP-a progenitor was retained at the site of origin of the AFP, as both genes were duplicated, in situ, multiple times (Fig. 2A). In sculpin, it was the 15-exon lunapark gene that gave rise to the AFP (Fig. 8C), but here, only small portions of the original gene were retained in the AFP. In contrast, the snailfish *AFP* does not share any similarity to functional genes. Instead, it likely arose from transposons and repetitive DNA (Fig. 6).

**Figure 8:**
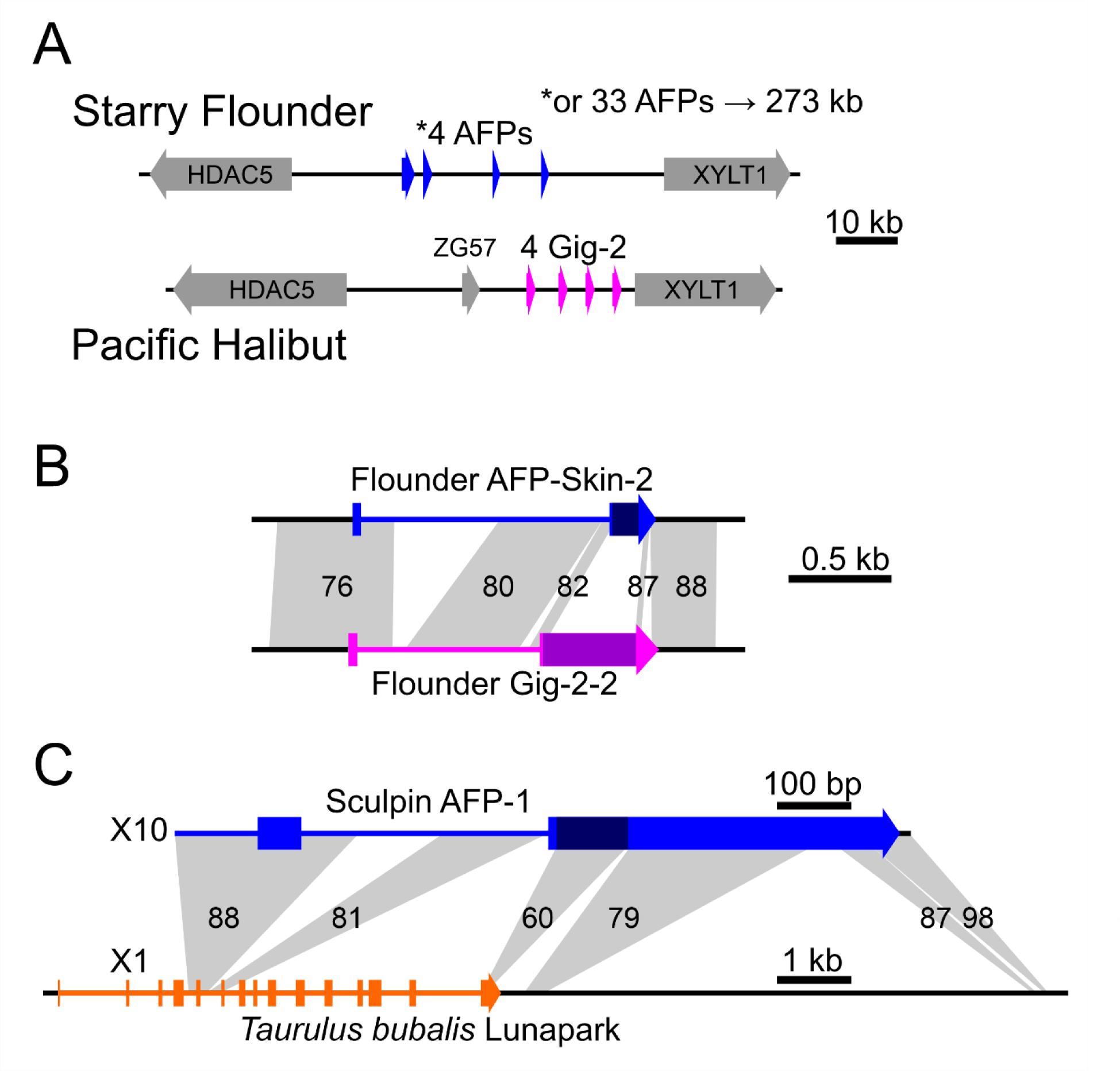
Origin of flounder and sculpin type I AFPs from progenitor genes. A) Comparison of the corresponding genomic regions, containing the HDAC5 and XYLT1 genes, of starry flounder and Pacific halibut. The two alleles from an individual flounder contained either 4 or 33 AFP gene copies. B) Comparison of a single flounder Gig-2 and skin AFP gene. The percent identity between the homologous regions (grey shading) is indicated, with the two exons (wide) and intron (narrow) of each gene in blue or purple. The portion of the second exon containing the coding region is in a darker shade. C) Comparison of the AFP gene of sculpin with the lunapark locus of a closely related fish, denoted as above, except the lunapark exons are in orange and lunapark is scaled 1:10 relative to the smaller AFP gene.

The Ala-rich coding sequences of the *AFP*s are the portions of the genes that have diverged the most since their origins, consistent with positive selection for a new function. Still, there are clues as to their origins, and these also differ between the four groups. In flounder, a very small helical region of Gig-2, encoding a single Thr and three Ala, was likely tandemly amplified many times while the rest of the coding sequence was lost. As two of the three Ala were encoded by GCC in the Gig-2 gene (40), this may explain the preference for GCC in the AFPs of this species (38). In sculpin, the coding sequence arose via frameshifting and mutation of a Glu/Gln-rich region at the end of the GIMAP-a coding sequence, giving rise to an Ala-rich region with a preference for GCG (substitution within Gln (GAG) codon) and GCA (frameshifted Glu (CAG) codon) (42). However, in cunner, a long Ala-rich stretch was present in the progenitor prior to the origin of the AFP (Fig. 4), and the GCT bias present in the GIMAP gene was retained in the AFP genes. The strong bias towards GCC in the snailfish AFP (Fig. 6) could be due to conversion of repetitive DNA (GCC)_n_ into AFP coding sequence.

## Discussion

AFPs of diverse structure are found in a wide variety of organisms (2). β-helical folds are prominent in all but fish and have arisen via convergence multiple times. In contrast, the helical type I AFPs have only arisen in fish, albeit four times. Most type I AFPs, as well as the other types of fish AFPs, are only moderately active (58). The reason for this may be related to the different environments that AFP-producing species inhabit. The oceans do not cool below ∼-1.8 °C, so fish only require ∼1 °C of TH protection together with ∼0.8 °C of colligative freezing point depression(5, 6), whereas terrestrial organisms can be exposed to much colder temperatures. β-helical AFPs can have large ice-binding surfaces (59, 60), and small increases in the area of the ice-binding face, through duplication of coils, can result in large increases in activity (61, 62). A single straight α-helix can only generate a narrow ice-binding face, and the activity increase upon duplication of a repeat is more modest (63). Therefore, these moderately active AFPs types are unlikely to be of much use to terrestrial organisms, whereas they are active enough to protect fish. That being said, type I AFPs with significantly higher activity have been found that either dimerize (35) or form multimers (64), but these appear to have arisen from much shorter progenitors (40).

The TH activity of cunner plasma during the winter is low (0 to 0.16 °C) (29, 65), with higher activity in the skin that is still below 1 °C (65), even in fish living in the ice-laden waters off Newfoundland. This modest activity can be explained by two factors. First, all of the isoforms are expected to fold as isolated α-helices, albeit with longer isoforms that are likely to be somewhat more active. Second, there are only eleven AFP genes within the fish that was sequenced, which came from New Brunswick waters (GenBank BioSample SAMN22589422). Ocean pouts (type III AFP) from the same waters have around 40 gene copies (66). However, cunner become torpid and do not feed during the winter (65), so they may not require high levels of AFPs throughout their bodies, in contrast to more winter-active species.

Snailfishes have much higher TH levels in their plasma than cunner, averaging 0.73 °C in the Atlantic snailfish and 0.92 °C in the dusky snailfish (57). Both of these fishes inhabit colder waters than Tanaka’s snailfish (67). The fish used to generate the chromosomal-level genome assembly for which only one AFP gene was found (Tanaka-6) was caught in the Yellow Sea (53). The other fish, with at least eight gene copies, originated from the Sea of Japan, a much icier location (68). As multigene families are notoriously difficult to assemble from Illumina reads, this may be an underestimate of the true gene copy number. These AFPs are significantly longer (81 – 137 a.a.) than those of the cunner (47 – 76 a.a.) and rather than forming extended α-helices, they are predicted by AlphaFold2 to form hairpins that likely dimerize in antiparallel fashion, similar to the larger (195-a.a.) dimeric isoform of winter flounder (35). If so, two species will have independently generated Ala-rich AFPs that are more complex than a single extended α-helix that fold in the same way.

When Evans and Fletcher probed a snailfish cDNA library using AFP sequences from sculpin and flounder, they did not obtain AFP sequences. Instead, they recovered clones encoding keratin or eggshell proteins that could potentially encode an Ala-rich protein, either directly or via frameshifting (54). While this was a solid hypothesis, the AFP genes herein do not share any similarity with these genes outside of the coding sequence. Interestingly, frameshifting did appear to play a role in the origin of the sculpin AFP from the lunapark locus (42). Without the presence of conserved flanking non-coding DNA, the similarity between *lunapark* and the *AFP* would have been unrecognizable as the AFP coding sequences are under strong selection for their new function. With nothing but TEs flanking the snailfish genes, which are present hundreds of times with the genome, the progenitor of the AFP coding sequence of snailfishes remains unverified. However, the codon bias is consistent with an origin from DNA rich in GCC repeats.

The three genes that gave rise to type I AFPs have different functions. The GIMAP family of proteins, one of which gave rise to the cunner AFP, are generally found in the cytoplasm of lymphocytes, where their loss has been shown to be detrimental to cellular function in mammals. They have an N-terminal GTP-binding domain and C-terminal tails of different lengths that likely confer unique properties, such as membrane anchoring, to the different family members (69). The Gig2 proteins that gave rise to the flounder AFP also belong to a family that is involved in immune function, but this proteins appears to be restricted to non-amniote vertebrates, with overexpression leading to resistance to viral infection (70). The progenitor of the sculpin AFP, lunapark, is not a member of a multigene family, nor is it involved in immune function. Instead, this protein is involved in the modelling of junctions of the endoplasmic reticulum (43). Other fish AFPs have also arisen by neofunctionalization of duplicated genes with unrelated functions. Type II arose from C-type lectins (11), type III from the C-terminal domain of sialic synthase (7), and the AFGPs of Antarctic fishes from trypsinogen (20).

It is not just pre-existing genes that have given rise to AF(G)Ps as the AFGP of northern cods arose from non-coding DNA (22). The snailfish gene may also have arisen from repetitive non-coding DNA, as well as TEs that define its non-coding regions. Despite the myriad detrimental effects that TEs have on the genome, they also have positive effects and their regulatory sequences have been co-opted by a number of genes (71, 72). In addition, numerous genes have arisen through ‘domestication’ of TEs, including proteins involved in the function of the brain and placenta (72), as well as a large number of transcription factors (73). They can also act as drivers of gene duplication, even after they have lost the ability to transpose on their own (72).

Positive selection has acted at two levels to enable AFP producing fish to avoid freezing. Firstly, the similarities between the type I *AFPs* of sculpins (42), cunner, and flounder (40) and their progenitor genes are highest between non-coding sequences, and much lower or barely detectible between the coding sequences. These are extreme examples of the positive selection that can drive diversification of duplicated genes (74). Secondly, all of these genes are present in multiple copies, indicative of positive selection for increased dosage in response to environmental stress (75). An additional benefit is that with more gene copies, there is a greater chance that one of them will accrue a beneficial mutation. Amplification of the beneficial variant will again generate a greater number of targets for additional beneficial mutations (76), resulting in rapid evolution of the gene family through multiple rounds of birth-and-death evolution (77). Positive selection of gene duplicates can usually be measured by comparing the ratio of non-synonymous (change in amino acid) to synonymous (silent) mutations in the copies. Proteins under strong positive selection for a new function will have ratios near to or exceeding unity, whereas those under negative selection have ratios well below one (75). In the case of these AFPs however, the ratio cannot be calculated as the coding sequences have diverged to such a degree that they cannot be accurately aligned, making a one-to-one correlation of their codons impossible. Nevertheless, these sequences are/were clearly subject to strong positive selection as the flanking sequences can be accurately aligned as they have diverged at a far slower rate.

In summary, the convergent evolution of AFPs in numerous lineages, in which multiple AFP types were generated from different progenitors, suggests that the onset of global cooling around 30 MA (78) was a strong environmental stressor for fish. It is remarkable that the Ala-rich α-helix type I AFPs arose independently in four different fish lineages, but not in any other organisms. Equally remarkable are the varied mechanisms by which similar AFPs arose via convergent evolution from both duplicated genes and from non-genic sequences. The common thread is that, at least in the case of the AFPs derived from pre-existing genes, the initial gene duplication event was followed by rapid diversification and additional gene duplication events.

## Supporting information

Supplementary figures and tables for Graham and Davies

## Acknowledgements

We are grateful for the pioneering work done by Drs. Rod Hobbs and Garth Fletcher to characterize the cunner and snailfish AFPs. This research was supported by CIHR Foundation Grant FRN 148422 and NSERC Discovery Grant RGPIN-2016-04810 to PLD.

## Notes

### Competing Interest Statement

The authors have declared no competing interest.

