## Supplementary figures and tables for Graham and Davies for "Convergent evolution of type I antifreeze proteins from four different progenitors in response to global cooling"

Included herein are figures S1 to S3 and tables S1 to S4

```
cunner-a1 QQDLRIVLLGKTGSGKSSTGNTILG-REAFRAKFSPSSVTNTCKKVFFQYEDRIVSLIDTPGIFDTSITEVKLKEEIVKY 79
cunner-a2 QQDLRIVLLGKTGSGKSSTGNTIVG-REAFRAKFSPSSVTNTCKKVFFQYEDRIVSLIDTPGIFDTSITEVKLKEEIVKY 79
cunner-a3 QQDLRIVLLGKTGSGKSSTGNTIVG-REAFRAKFSPSSVTKTCKKVFFQYEDRIVSLIDTPGIFDTSITEVKLKEEIVKY 79
cunner-a4 QQDLRIVLLGKTGSGKSSTGNTILG-REAFRAKFSPSSVTNTCKKVFFQYEDRIVSLIDTPGIFDTSITEVKLKEEIVKY 79
cunner-a5 QEDLRIVLLGKTGSGKSSTGNTILG-RKAFEAEFSPSSVTKTCRKVLFQYKDRRVRLIDTPGIFDTSIKEVKLKEEIQKC 79
ballan-a1 QEDLRIVLLGKTGSGKSSTGNTILG-NEVFKAKCSASSVTRTCREVVLQYKDRRVRLIDTPGIFDTSIAEVKLKEEIETC 79
ballan-a2 QEDLRIVLLGKTGSGKSSTGNTILG-NEVFKAEFSASSVTKTCQKVFFHYKDRGVSLIDTPGIFDTSITEVELK-EIEEC 78
ballan-a3 QEDLRIVLLGKTGSGKSSTGNTILG-NEVFNTDVSPSSVTKTCKKVFFYYKDRRMSVIDTPGIFDTSIAEVELKEEIEKC 79
ballan-a4 QEDLRIVLLGKTGSGKSSTGNTILG-REVFKAECSASSVTKTCRKVFFHYKDRRVRLIDTPGIFDTSIAEVQLKEEIEKC 79
spotty-a  TKDLRIVLLGKTGSGKSSTGNTIAG-REVFKAGVSPSSLTKTCTKVMSHHDGRSVSIIDTPGVFDTSTTNEELKDEIEKC 79
spotty-b  SGGLRIVLVGKTGSGKSATGNTILG-RAAFKEDPSPVSVTKHCETQNGEVDGTLVQVIDTPGLFDTGITEEELKTRIEEC 79
ballan-b1 SSGLRIVLVGKTGSGKSATGNTILG-RAAFKEDPSSISVTKNCATQSGEVDGTAVQVIDTPGLFDTDITEKELKTRIEEC 79
ballan-b2 PGGLRIVLVGKTGSGKSATGNTILG-RAAFKEDPSPVSVTKNCATQSGEVDGTAVQVIDTPGLFDTDITEKELKTRIEEC 79
cunner-b1 STGLRIVLVGKTGSGKSATGNTILG-RSVFKEDPSPVSVTKHCETQSGEVDGTAVQVIDTPGLFDTGITEEELKTRIEEC 79
cunner-b2 PTGLRIVLVGKTGSGKSATGNTILGVLDAFKEDPSPASVTKHCKNQSGEVDGTAVQVIDTPGLFDTDITEEELKTRIEEC 80
spotty-c1 QGDLRLILVGKTGSGKSASGNTILGYRYAYKTDISPESVTVGCHKEEVQDRGRNLFVIDTPGLFDTRKTTEGLKGDIEEC 80
spotty-c2 QGDLRLILVGKTGSGKSASGNTILGRRYAFETDISPESVTVGCHKEKVQDRGRNLLVIDTPGLFDTRKTTEDVKGDIEKC 80
spotty-c3 QGDLRLILVGKTGSGKSASGNTILGRRYAFETDISPESVTAGCHQEEVKDGERNLFVFDSPGLFDTSKTTEDVKEDIEDC 80
spotty-c4 QGHLRLILVGKTGSGKSASGNTILGRRYAFETDISPESVTAGCHQEEVQDGERNLFVIDSPGLFDTSKTTEDVKEDIEDC 80

cunner-a1 ILLSVPGPHVFLLVIKLD-RFTEKEKNAVKWIEDNFGAEASKYTLVLFTRGDELGQKSIETFLVESSDLRKFLVA 153
cunner-a2 ILLSVPGPHVFLLVIKLD-RFTEKEKNAVKWIEDNFGAEASKYTLVLFTRGDELGQKSIETFLVESSDLRKFLVA 153
cunner-a3 ILLSVPGPHVFLIVIKLD-RFTEKEKNAVKWIEDNFGAEASKYTLVLFTRGDELGQKSIETFLVESSDLRKFLVA 153
cunner-a4 ILLSVPGPHVFLLVIKLD-RFTEKEKNAVKWIEDNFGAEASKYTLVLFTRGDELGQKSIETFLVESSDLRKFLVA 153
cunner-a5 IHLSVPGPHVFLLVIRLDVRFTEEEKNAFKWIDDNFGAEASKFTLVLFTRGDELGQKSIETFLDTSSDLRDLIRT 154
ballan-a1 ILLSVPGPHVFLLVIRLDVRFTQEEKNVVKWIKDNFGAEASKYTLVLFTRGDELGQKSIETFLEDSSDLKELIRT 154
ballan-a2 ILLSVPGPHVFLLVIRLDVRFTENEKNVVKWIKDNFGAEASKYTLVLFTREDQLGQKSIETFLEESSDLKELIRT 153
ballan-a3 ILLSVPGPHVFLLVIRLDVRFTQEEKNTVKWIKDNFGAEASKYTLVLFTREDELGQKSIETFLEESSDLRDLIRM 154
ballan-a4 ILLSVPGPHVFLLVIRLDVRFTENEKNAVKWIKDNFGAEASKYTLVLFTRGDQLGQKSIETFLEESADLKELIRT 154
spotty-a  IMMSLPGPHVFLLVISLDARFTEEEKKAVTWIKDNFGEEASKYTLVLFTRGDVLRETSIQTFLKESPELQEVIRT 154
spotty-b  VKMSVPGPHAFLLVIRLGVRFTEEERNAVKWIQDNFGDDASMYTIMLFTCKD---QAKADNALKECKELRRLSIT 151
ballan-b1 VEMSVPGPHAFLLVIRLGVRFTEEERNAVKWIQDNFGEDASMYTIMLFTCKD---QAKADNALKECKELRRLSIT 151
ballan-b2 VEMSVPGPHAFLLVIRLGVRFTEEERNAVKWIQDNFGEDASMYTIMLFTFKD---QAKAEDALKECKELRKIQDS 151
cunner-b1 VKMSVPGPHAFLLVIRLGVRFTEEERNAVKWIQDNFGDDASMYTIMLFTCKD---QAKADNALKECKELRRLSIT 151
cunner-b2 VKMSVPGPHAFLLVIRLGVRFTAEERNAVKWIQDNFGDDASMYTIVLFTFKD---QANVDDALKDCKELRKITDS 152
spotty-c1 VAQSVPGPHAFLLVISLKARFTEEEKASVKWIQDNFGPKSSLYTMILFTHADLLEGKTVEDFVRESKHLQKLITE 155
spotty-c2 VVQSVPGPHAFLLVISLKARFTEEEKASVKWIQDNFGPESSMYTMILFTHADLL--------------------- 134
spotty-c3 IVQSVPGPHAFLLVISLKAKLTEQEKASVKWIQDNFGPKSSMYTMILFTHADLLGEKTVEDFVRESKHLQKLINE 155
spotty-c4 VVQSVPGPHAFLLVISLKARFTEEEKASVKWIQDNFGPKSSMYTMILFTHADLLGEKTVEDFVRESKHLQKLITQ 155
```

**Supplementary Figure 1**: Alignment of the GTPase domain of the GIMAP isoforms from cunner, ballan wrasse, and spotty, coloured using 'Color Align Properties' from the Sequence Manipulation Suite (1). This alignment was used to generate the phylogenetic tree shown in Fig. 3.

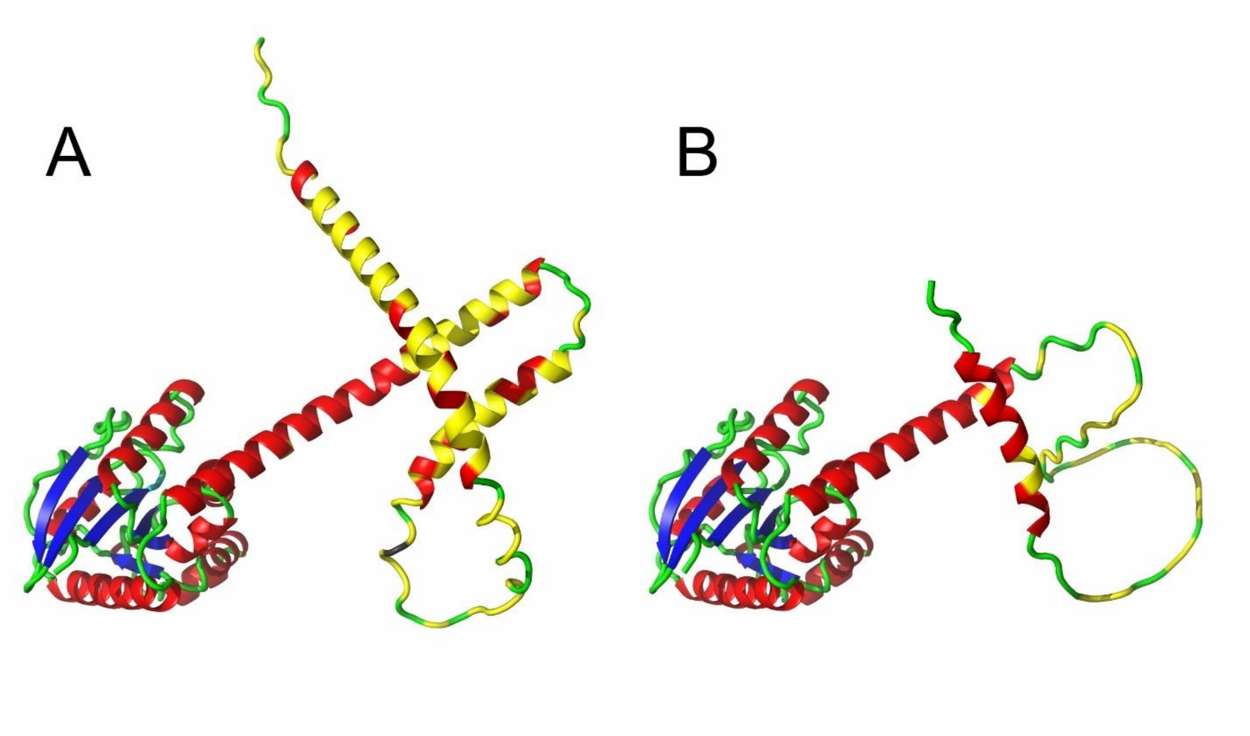


**Supplementary Figure 2**: AlphaFold2 prediction of the structures of A) Ballan wrasse GIMAP-a1 and B) Cunner GIMAP-a5 rendered using cartoon mode in PyMOL (2). The GTPase domain (globular domain at left) was predicted with high confidence while the prediction of the C-terminal extension was with low confidence. Helices are red, sheets cyan, and loops green, with Ala residues in the extension in yellow.

```
tggggtccgc tggacagttt ataggggcgc acccggtggg gaggcggggt agacatctgt   60
cctgtgtctc tgccggcgtg agggaaacgt attccgtcct gagtttccac actcctagct  120
ctccaaacta tctctctctc taaactctca gccaacaacc cacattttga atggagcctg  180
aggggtttat attgctgcca aacctcccgg caatctgaac cgcttgccga agcagtcacg  240
tggtacagcc tggcagcgag gcttcggatg tcctcatttt tggctcctcc cctaaatgaa  300
gcaagcctcg atacgcactt cgaggaaacg cccgctccat tactcgacac ttgcttataa  360
                                                        M  A  P
aagtcggcac atctcgtcac aacactaaca gaaggagaaa cagaaggatc atggcccctg  420
A  T  P  A   Q  R  A   A  A  T   A  A  D  A   A  A  K   A  A  F
ctacccccgc ccagagagcc gctgccaccg ccgctgacgc cgccgccaaa gccgctttca  480
T  A  A  T   V  A  A   G  V  A   D  T  A  A   K  V  S   E  A  S
ccgctgctac cgtcgccgct ggcgtcgccg acaccgccgc caaagtctcg gaagcctccg  540
A  A  T  V   A  K  T   V  V  A   A  K  A  A   A  T  A   V  A  P
ctgccaccgt cgctaaaact gtcgtcgccg ccaaagccgc tgccaccgcc gttgccccca  600
N  T  G  A   I  T  A   A  A  A   A  S  A  T   A  D  A   A  A  K
acaccggtgc catcaccgcc gccgccgctg cctctgccac cgccgacgcc gccgccaaag  660
A  A  R  A   T  A  D   A  A  A   A  K  A  A   A  A  G   V  T  A
ccgccagagc caccgccgac gccgctgccg ccaaagccgc cgctgccggt gtcaccgcca  720
K  A  A  A   A  A  L   A  A  L   *
aagccgccgc cgctgccctc gccgcccttt aaagggaaac ccaaagcagg acattttatc  780
agtggcctca agtgagcttg ggtttagttg ggatcatgtg ttgtcctgta ttatgattat  840
aattcatatt attgttatta tttgtacctg tcacgcgcta taaaaggtga cgtgtcatgt  900
gtgttcattt tgggagcttg cagaagcctt gaatttgatt taataaaaca atattttgga  960
ttcccgcata tcgattatgt gtttttcttt cattaccatt taacgctcta caggttaagc 1020
aggaactgaa cagccattag cagatccgcc cattcccttt tttctttgcg tattgtagct 1080
ttatatatat atgaggttca tggttggtg                                  1109
```

**Supplementary Figure 3**: Sequence of the Dusky-3 AFP locus assembled from Illumina reads from SRA dataset SRX18366605. The translation of the intronless AFP coding sequence is shown above the corresponding sequence.

**Supplementary Table 1**: Verification of GIMAP annotation, reannotated sequences are below.

| Sequence | GenBank Accession | Status |
|---|---|---|
| Ballan GIMAP-a1 | XP_020509819.1 | Second intron unsupported |
| Ballan GIMAP-a2 | XP_029137074.1 | Second intron unsupported |
| Ballan GIMAP-a3 | XP_020509833.1 | Verified |
| Ballan GIMAP-a4 | XP_020509831.1 | Verified |
| Ballan GIMAP-b1 | XP_020509835.1 | Verified |
| Ballan GIMAP-b2 | XP_029137070.1 | Verified |
| Spotty GIMAP-a | XP_034543013.1 | Supported |
| Spotty GIMAP-b | XP_034544901.1 | Earlier start codon not consistent |
| Spotty GIMAP-c1 | LOC117816646 | Only isolated exons predicted, listed as pseudo |
| Spotty GIMAP-c2 | LOC117815504 | Only isolated exons predicted, listed as pseudo |
| Spotty GIMAP-c3 | LOC117815504 | Only isolated exons predicted, listed as pseudo |
| Spotty GIMAP-c4 | XP_034544162.1 | First exon not consistent |

```
>Ballan GIMAP-a1
MALRSTKVCGSPDIQEDLRIVLLGKTGSGKSSTGNTILGNEVFKAKCSASSVTRTCREVVLQYKDRRVRLIDTPGIFDTSIAEVKLKEEIETCILLS
VPGPHVFLLVIRLDVRFTQEEKNVVKWIKDNFGAEASKYTLVLFTRGDELGQKSIETFLEDSSDLKELIRTCNARYIVFDNKSMNNRTQVNDLFEKI
DEIVQLNGGHYTSSIYAEAQRKIKSAEWWNTCVQYVPLLSAGAAAAAAAAGAAAVAIGTGASAAAIGAGAAAAAAGAAAVAIGAGATAAADGAGAAA
AAVGVGAAAAAGIGAAAAAIGAGAAAAVGAAAAAAAAAAAAGAAAAIGAAAGPRAAG

>Ballan GIMAP-a2
MALRSTKVCGSPDIQEDLRIVLLGKTGSGKSSTGNTILGNEVFKAEFSASSVTKTCQKVFFHYKDRGVSLIDTPGIFDTSITEVELKEIEECILLSV
PGPHVFLLVIRLDVRFTENEKNVVKWIKDNFGAEASKYTLVLFTREDQLGQKSIETFLEESSDLKELIRTCNSRYIVFDNKSMNNRTQVNDLFEKID
EIVQLNGGHYTSSIYEEAQRKIKSDKRWNNFLIHQMSPLSTSMKP

>Spotty GIMAP-b
MAEAIPLREKKQKTLPRSDGCGLNNFDALDLSGGLRIVLVGKTGSGKSATGNTILGRAAFKEDPSPVSVTKHCETQNGEVDGTLVQVIDTPGLFDTG
ITEEELKTRIEECVKMSVPGPHAFLLVIRLGVRFTEEERNAVKWIQDNFGDDASMYTIMLFTCKDQAKADNALKECKELRRLSITFGRRYHAFNNND
ADDRVQVNELITMIKDMIQDNGGKHYTNEMYERAQRKLREEEERRKQEEEEKKEEERKVWDAEREKQEKERAKKKNVRIKNIRVASAAAAVLVVAGV
VIAVGASNTVALALGAPVLVLGVLCGLAAICVWKGIRCKAKKGVPV

>Spotty GIMAP-c1
MLRIKAWIPGCTLLVLVTLCGQTAHCQSQLKDSPHQGDLRLILVGKTGSGKSASGNTILGYRYAYKTDISPESVTVGCHKEEVQDRGRNLFVIDTPG
LFDTRKTTEGLKGDIEECVAQSVPGPHAFLLVISLKARFTEEEKASVKWIQDNFGPKSSLYTMILFTHADLLEGKTVEDFVRESKHLQKLITECGGP
IPLAHQ

>Spotty GIMAP-c2
MLRIKAWIPGCTLLVLVTLCGPTAHCQSQLKDSSHQGDLRLILVGKTGSGKSASGNTILGRRYAFETDISPESVTVGCHKEKVQDRGRNLLVIDTPG
LFDTRKTTEDVKGDIEKCVVQSVPGPHAFLLVISLKARFTEEEKASVKWIQDNFGPESSMYTMILFTHADLL

>Spotty GIMAP-c3
MLRIKAWIPGCTLLVLVTLCGPTAHCQSQLKDSTHQGDLRLILVGKTGSGKSASGNTILGRRYAFETDISPESVTAGCHQEEVKDGERNLFVFDSPG
LFDTSKTTEDVKEDIEDCIVQSVPGPHAFLLVISLKAKLTEQEKASVKWIQDNFGPKSSMYTMILFTHADLLGEKTVEDFVRESKHLQKLINECGGR
YHSLINDNKQSRDQVRQLIDKIDTMVESNGGSHYTSEMYQRAQEQVEEERRREEEQEKEEAKRAKCKAVMMALAGITGGSFLFPSYLFTLVAGGLGP
EALDCIKDMFK

>Spotty GIMAP-c4
MLRIKAWIPGCTLLVLVTLCGPTAHCQSQHKDSTHQGHLRLILVGKTGSGKSASGNTILGRRYAFETDISPESVTAGCHQEEVQDGERNLFVIDSPG
LFDTSKTTEDVKEDIEDCVVQSVPGPHAFLLVISLKARFTEEEKASVKWIQDNFGPKSSMYTMILFTHADLLGEKTVEDFVRESKHLQKLITQCGGR
YHSLINDNKQSRDQVRQLIDKIDTMVEFNGGSHYTSEMYQRAQEQIEEERRWEEEQEKEEAKRAKCKAVMVALAGIAGVSYLFPSYLLTLVAGTFGP
EAYDCIKYMFK
```

**Supplementary Table 2**: Presence or absence of type I AFP sequences in sequence databases from fishes in family Liparidae.

| Database Accession | Type | Species | Tissue | Collection Date | Location | Result |
|---|---|---|---|---|---|---|
| SRLO00000000 | Genome | *Liparis tanakae* | Muscle | 2017 | Korean Coast | +++ |
| SRR22396815 | SRA DNA | *L. gibbus* | | 2011 | Bering Sea | +++ |
| SRR12518887 | SRA DNA | *L. liparis* | | | North Sea | +++ |
| SRR22396704 | SRA DNA | *L. tunicatus* | | 2010 | Bering Sea | +++ |
| SRR19970642 | SRA RNA | *Pseudoliparis swirei* | Skin | | Mariana Trench | No obvious AFP |
| SRR19963796 | SRA DNA | *P. swirei* | | | Mariana Trench | No obvious AFP |
| RZNG00000000 | Genome | *Pseudoliparis* sp. | | | Yap Trench | No obvious AFP |
| SRR7206520 | SRA RNA | *P. amblystomopsis* | Liver | | Mariana Trench | No obvious AFP |
| SRR7223797 | SRA DNA | *P. amblystomopsis* | | | N. Pacific depths | No obvious AFP |
| SRR12575254 | SRA RNA | *P. amblystomopsis* | Skin | | Yap Trench | No obvious AFP |
| SRR21844227 | SRA DNA | *Paraliparis pectoralis* | | 1996 | Oregon Coast | No obvious AFP |
| SRR22396715 | SRA DNA | *Psednos* sp. | | 2015 | N. Carolina Coast | No obvious AFP |
| SRR22396812 | SRA DNA | *Careproctus phasma* | | 2011 | Bering Sea | No obvious AFP |
| SRR22396818 | SRA DNA | *Careproctus scottae* | | 2011 | Bering Sea | No obvious AFP |
| SRR22396822 | SRA DNA | *Crystallichthys cyclospilus* | | 2010 | Aleutian Islands | No obvious AFP |

**Supplementary Table 3**: Characteristics of scaffolds/chromosomes from Tanaka's snailfish that contain AFP sequences and similarity to RNAseq reads within dataset SRR8617419.

| **Accession** | **Scaffold** | **Length (kb)** | **Coding sequence** | **Notes** |
|---|---|---|---|---|
| SRLO01021987 | 21988 | 1.38 | Tanaka-3 | 96% match to RNA |
| SRLO01001937 | 1938 | 25.03 | Tanaka-2 | 99% match to RNA, downstream Ala-rich x2, limited RNA support |
| SRLO01012854 | 12855 | 2.31 | Tanaka-1 | ~89% match to RNA over entire length |
| SRLO01024077 | 24078 | 1.23 | Tanaka-4 | One highly similar RNA fragment (97%) |
| SRLO01015817 | 15818 | 1.95 | Tanaka-5 | 99% match to RNA |
| SRLO01010016 | 10017 | 2.76 | Partial | Coding sequence contains a gap |
| SRLO01021795 | 21796 | 1.40 | Partial | Scaffold begins mid-AFP |
| SRLO01013587 | 13588 | 2.21 | Partial | N-terminus missing within a gap |
| CM070098 | Chr 11 | 22921 | Tanaka-6 | Coding starts at base 3855161<br>99% match to RNA |

**Supplementary Table 4**: Characteristics of transposable element sequences flanking Tanaka's snailfish AFP sequences.

| Transposon | Censor match | Best Score |
|---|---|---|
| TE-1 | hAT-N5_CCa (carp) | 442 |
| TE-2 | hAT-4_Ilyo (eel) | 1710 |
| TE-3 | Copia-1_PeFla-I (perch) | 1205 |
| TE-4 | hAT-36_Ilyo (eel) | 1059 |
